# The respiratory microbiome and susceptibility to influenza virus infection

**DOI:** 10.1101/372649

**Authors:** Kyu Han Lee, Aubree Gordon, Kerby Shedden, Guillermina Kuan, Sophia Ng, Angel Balmaseda, Betsy Foxman

## Abstract

Influenza is a major cause of morbidity and mortality worldwide. However, vaccine effectiveness has been low to moderate in recent years and vaccine coverage remains low, especially in low- and middle-income countries. Supplementary methods of prevention should be explored to reduce the high burden of influenza.

A potential target is the respiratory tract microbiome, complex microbial communities which envelop the respiratory epithelium and play an important role in shaping host immunity. Using a household transmission study, we examined whether the nose/throat microbiota was associated with influenza susceptibility among participants exposed to influenza virus in the household. Further, we characterized changes in the nose/throat microbiota to explore whether community stability was influenced by influenza virus infection.

Using a generalized linear mixed effects model, we found a bacterial community type associated with decreased susceptibility to influenza. The community type was rare and transitory among young children but a prevalent and stable community type among adults. Using boosting and linear mixed effects models, we found associations between the nose/throat microbiota and influenza also existed at the taxa level, specifically with the relative abundance of *Alloprevotella, Prevotella,* and *Bacteroides* oligotypes.

We found high rates of change in the bacterial community among both secondary cases and household contacts who were not infected during follow up. Preliminary results suggest short-term changes in the bacterial community structure may differ between the two groups, but further work is needed to validate our observations.

Lastly, age was strongly associated with susceptibility to influenza and the nose/throat bacterial community structure. Although additional studies are needed to determine causality, our results suggest the nose/throat microbiome may be a potential target for reducing the burden of influenza.

**Author summary:** Microbiome research has transformed our understanding of microbes and human health. Resident bacteria can protect the host from pathogens by shaping immunological responses. These new insights suggest the microbiome could be a target for preventing influenza virus infection, a major cause of illness and death worldwide. In this study, we explored the relationship between the nose/throat microbiota and influenza virus in Nicaraguan households.

Household members were enrolled immediately after one member was diagnosed with influenza virus infection. This study design allowed us to identify associations between the microbiota and influenza susceptibility. We also explored whether influenza virus infection altered the bacterial community structure and found short-term changes were common among both secondary cases and household members who remained influenza negative during follow up. Lastly, we found age played major roles in both influenza susceptibility and in short-terms changes in the microbiota. Although much work is needed to determine causal relationships, our findings suggest strategies that appropriately modify the microbiome might be useful in preventing influenza virus infections.

## Introduction

Influenza is a major contributor of human illness and death worldwide, estimated to cause 3-5 million cases of severe illness [1] and 400,000 deaths during interpandemic years [2]. Vaccination is the best available means of influenza prevention. However, vaccine effectiveness has been low to moderate in recent years [3,4] and vaccine coverage remains low, especially in low- and middle-income countries [5]. With increasing support for a role of the microbiome in shaping host immunity [6–8], exploring whether these effects extend to influenza risk could contribute to supplementary methods of prevention.

We hypothesized that the nose/throat microbiome is an unrecognized factor associated with susceptibility to influenza virus. Murine and human studies support this assertion. Compared to controls, mice treated with oral antibiotics exhibited enhanced degeneration of the bronchiole epithelial layer and increased risk of death following intranasal infection with influenza virus [7]. In two separate randomized controlled trials, newborns fed prebiotics and probiotics had significantly lower incidence of respiratory tract infections compared to placebo [9,10]. These studies suggest the manipulation of the microbiome, either through disruption or supplementation, can alter risk of respiratory tract infections.

The epithelial cells of the upper and lower respiratory tracts are the primary targets for influenza virus infection and replication [11]. However, these cells are enveloped by complex bacterial communities that may directly or indirectly interact with influenza virus to mediate risk of infection. Commensal bacteria may prevent infection by regulating innate and adaptive host immune responses [6,7]. In addition, this immune response might stimulate changes in the microbiome [12–14]. In a human experimental trial, young adults given intranasal administration of live attenuated influenza vaccine were characterized by increased taxa richness relative to the control group [15].

Further, influenza-related changes in the bacterial community structure might explain the enhanced risk of bacterial pneumonia and otitis media following influenza virus infection [16–19]. The most commonly detected causative organisms of bacterial pneumonia and otitis media increase in abundance in the upper respiratory tract following respiratory virus infection [20,21]. We previously showed that adults in the US with influenza virus infection expressed increased nose/throat carriage of *Streptococcus pneumoniae* and *Staphylococcus aureus* [20]. Similarly, other studies have observed an increase in pneumococcal density following rhinovirus infection [21] and changes in the microbiota during rhinovirus and respiratory syncytial virus infections [22]. Increased carriage elevates risk of invasive disease [23,24], potentially through more frequent microaspiration into the lung or migration to the middle ear [25]. However, an association between the nose/throat microbiome and influenza risk has not been demonstrated in human populations.

In this study, we used data from a longitudinal household transmission study of influenza to assess the relationship between the nose/throat microbiota and susceptibility to influenza virus infection and to determine whether influenza virus infection alters the bacterial community structure using an untargeted 16S rRNA taxonomic screen (Fig 1).

**Fig 1.**
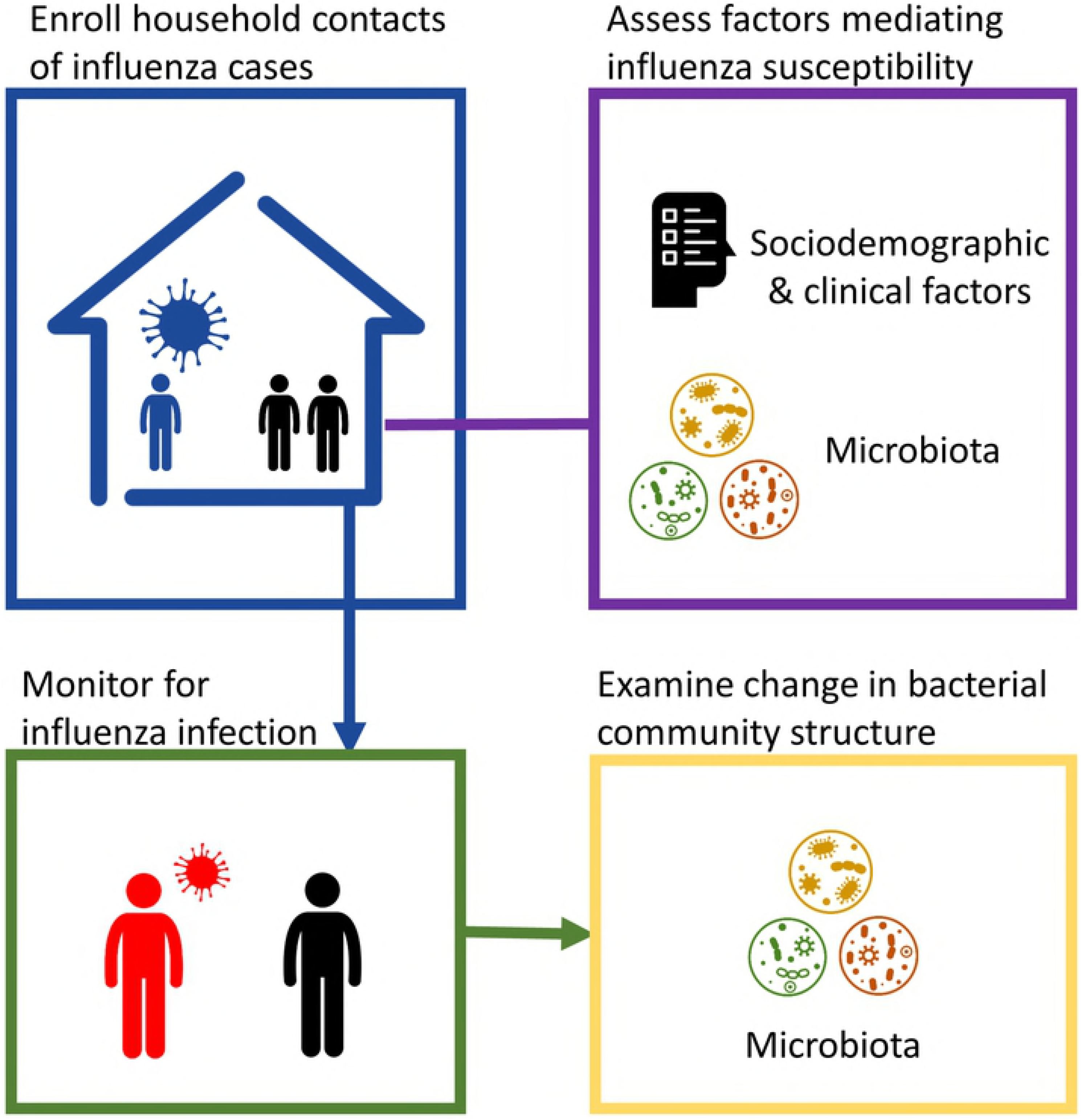
Graphical abstract.

## Results

### Study population

A total of 717 participants from 144 households were enrolled in the Nicaraguan Household Transmission Study during 2012-2014. During this period, 3,101 nose/throat samples were collected over 5 home visits (mean: 4.3 samples per person; interquartile range (IQR): 4-5). Analysis was restricted to 537 household contacts who were negative for influenza virus by real-time reverse transcription polymerase chain reaction (RT-PCR) at time of enrollment.

Sixty-one household contacts were children ≤5 years of age (median: 2 years; IQR: 1-4), 163 were children 6-17 years of age (median: 10 years; IQR: 8-14), and 313 were adults (median: 33 years; IQR: 24-43) (Table 1). Fifty-one percent of all household contacts were exposed to at least one tobacco smoker in the household and 29% resided in crowded households (on average, >3 persons per bedroom). Household contacts were rarely vaccinated against influenza (5%) and very few used antibiotics (<1% two weeks prior to enrollment and <1% during follow up) or oseltamivir (6% during follow up).

**Table 1.** Characteristics of 537 household contacts of influenza cases from 144 households, Managua, Nicaragua, 2012-2014, by community type* at enrollment.

| Characteristic | All<br>(n=537<sup>†</sup>) | Community<br>Type 1<br>(n=132) | Community<br>Type 2<br>(n=125) | Community<br>Type 3<br>(n=122) | Community<br>Type 4<br>(n=85) | Community<br>Type 5<br>(n=59) |
| --- | --- | --- | --- | --- | --- | --- |
|  | No. (%) | No. (%) | No. (%) | No. (%) | No. (%) | No. (%) |
| Age (years) |  |  |  |  |  |  |
| 0-5 | 61 (11) | 9 (7) | 15 (12) | 4 (3) | 3 (4) | 26 (44) |
| 6-17 | 163 (30) | 51 (39) | 41 (33) | 38 (31) | 19 (22) | 9 (15) |
| ≥18 | 313 (58) | 72 (55) | 69 (55) | 80 (66) | 63 (74) | 24 (41) |
| Female | 347 (65) | 86 (65) | 88 (70) | 87 (71) | 45 (53) | 35 (59) |
| Influenza infection | 71 (13) | 21 (16) | 20 (16) | 15 (12) | 5 (6) | 6 (10) |
| Influenza vaccination<sup>‡</sup> | 27 (5) | 8 (6) | 6 (5) | 8 (7) | 2 (3) | 3 (5) |
| Smoker in household | 245 (51) | 54 (46) | 58 (52) | 56 (51) | 44 (59) | 28 (50) |
| >3 persons per bedroom in the household | 156 (29) | 40 (30) | 35 (28) | 36 (30) | 22 (26) | 19 (32) |
| Antibiotic use <2 weeks prior | 1 (0) | 0 (0) | 0 (0) | 0 (0) | 0 (0) | 1 (2) |
| Antibiotic use during follow up | 4 (1) | 2 (2) | 0 (0) | 1 (1) | 0 (0) | 1 (2) |
| Oseltamivir use during follow up | 33 (6) | 4 (3) | 14 (11) | 6 (5) | 3 (4) | 5 (8) |

\*Community types were defined using Dirichlet multinomial mixture method (see Methods).
<sup>†</sup>Includes 10 household contacts with undefined community type at time of enrollment
<sup>‡</sup>Prior to enrollment and in same year as index case

After the enrollment of an index case, households were followed for up to 13 days through 5 home visits conducted at 2-3 day intervals. Seventy-one secondary cases from 48 households were identified by RT-PCR during follow up. Fourteen out of the 48 households had more than one secondary case (29%), suggesting clustering of cases by household. Most secondary cases were older children and young adults (median: 13.0 years; IQR: 6, 23) and had at least one symptom of an acute respiratory infection during follow up (79%) (S1 Table).

### Bacterial community types

We conducted 16S (V4) rRNA sequencing on a pair of samples from each study participant: 712 samples collected at enrollment and 698 samples collected at the last available home visit. The median time between samples was 9 days (IQR: 9-10 days). After quality filtering, microbiota data was available for 710 samples collected at enrollment and 695 samples collected at the last available home visit.

Dirichlet multinomial mixture modeling [26], an unsupervised clustering method, was used to assign nose/throat samples to 5 bacterial community types (S1 Fig: model fit by Dirichlet components; S2 Fig: PCoA plot by community type). Ninety-eight percent of all sequenced samples were assigned to a community type, after applying a ≥80% posterior probability threshold. Permutational multivariate analysis of variance (PERMANOVA) indicated community types differed significantly from one another (Bray-Curtis dissimilarity, p=0.001, R^2^=0.21). Relatively few oligotypes explained clustering of the single-community type model for the five-community type model, as 50% of the difference between models was attributed to 15 of the 230 total oligotypes. The relative abundances of these 15 oligotypes are depicted in Fig 2. The relative abundances of all 230 oligotypes are available in S2 Table.

**Fig 2.**
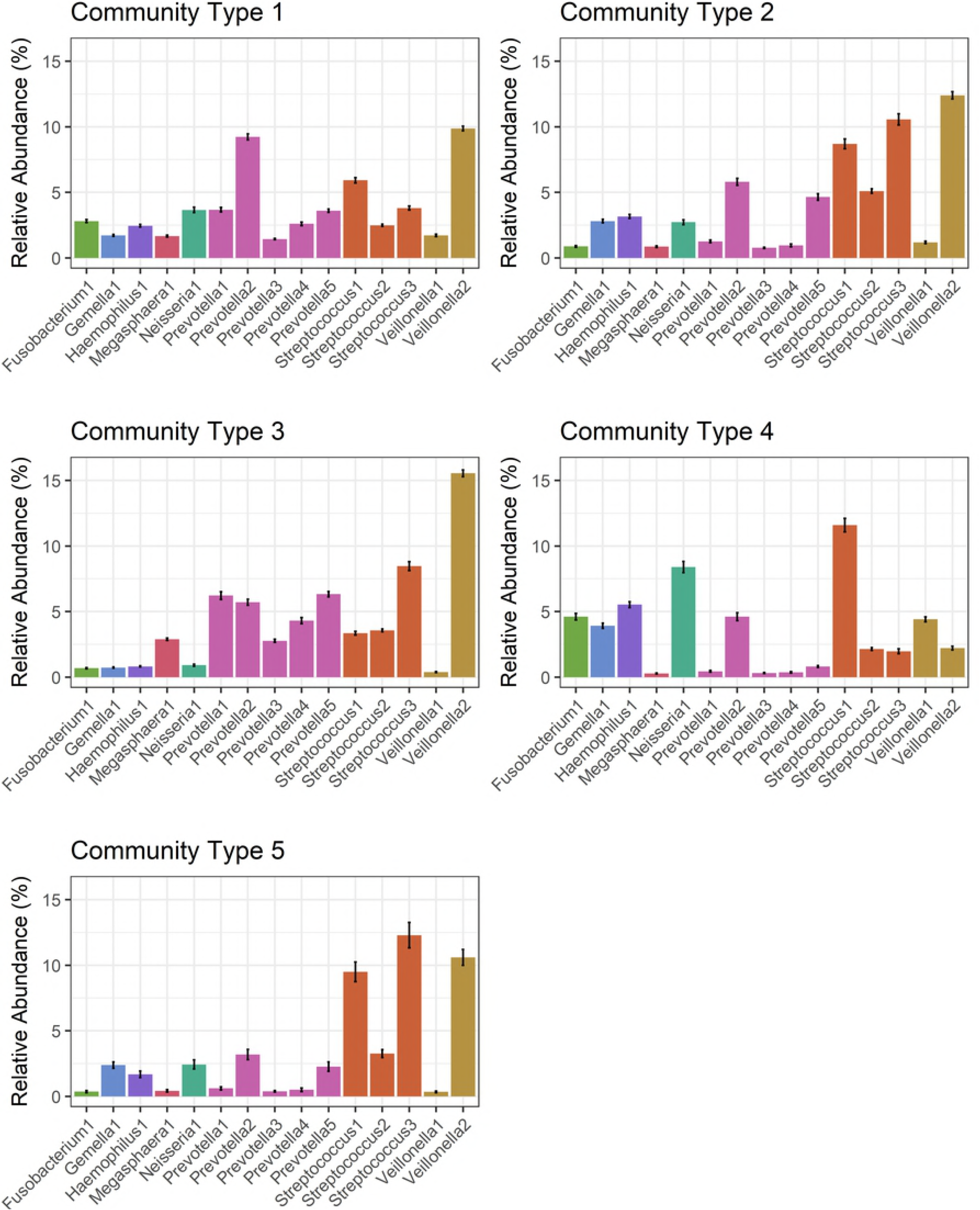
Relative abundance of 15 oligotypes, by community type. Oligotypes included in the figure attributed >50% of the difference between the single-community type model and the five-community type model. Bars represent the mean relative abundance of each oligotype (±1 standard error). 1,405 samples from 717 study participants residing in 144 households in Managua, Nicaragua, 2012-2014.

The prevalence of community types among household contacts differed significantly by age. Most notably, community type 4 was rare among young children and became more prevalent with age (at enrollment; 0-5 years: 5%, 6-17 year: 12%, adults: 20%; χ2-test, p=0.004) (Table 1). We observed similar results after restricting our analysis to household contacts who remained influenza negative during follow up (at enrollment; 0-5 years: 7%, 6-17 years: 12%, adults: 21%; χ2 test, p=0.011) (S3 Table). Young children were primarily colonized by community type 5, which was less common among older age groups (at enrollment 0-5 years: 43%, 6-17 years: 6%, adults: 8%; χ2-test, p<0.001) (Table 1). These results indicate age is strongly associated with the nose/throat bacterial community structure.

### Bacterial community type associated with lower susceptibility to influenza virus infection

To investigate the relationship between bacterial community types and influenza susceptibility, we first estimated secondary attack rates by community type. Secondary attack rates were calculated as: the number of secondary cases identified by RT-PCR during follow up over the total number of household contacts at start of follow up. Secondary attack rates among household contacts with community type 4 were nearly half of other community types; however, differences were not statistically significant (5.9% vs. 10.2%-16.0%; χ^2^-test, p=0.056) (Fig 3). Similar differences were observed after stratifying by age.

We used a generalized linear mixed effects model to examine the relationship between community types and influenza susceptibility after adjusting for age, a smoker in the household, household crowding, and clustering by household. A detailed description of the model is available in S1 Appendix. Our decision to account for household clustering was supported by an intra-class correlation of 0.21, which indicates 21% of the total variance was due to clustering by household. We found household contacts with community type 4 had a lower odds of influenza virus infection (odds ratio (OR): 0.26; 95% CI: 0.07, 0.99) (Fig 4), Further, young children were most likely to acquire influenza virus (OR: 4.66; 95% CI: 1.62, 13.37), followed by older children (OR: 2.91; 95% CI: 1.47, 5.80). These results suggest household contacts with community type 4 were less likely to be infected with influenza and younger household contacts were at greater risk after adjusting for other known risk factors.

**Fig 3.**
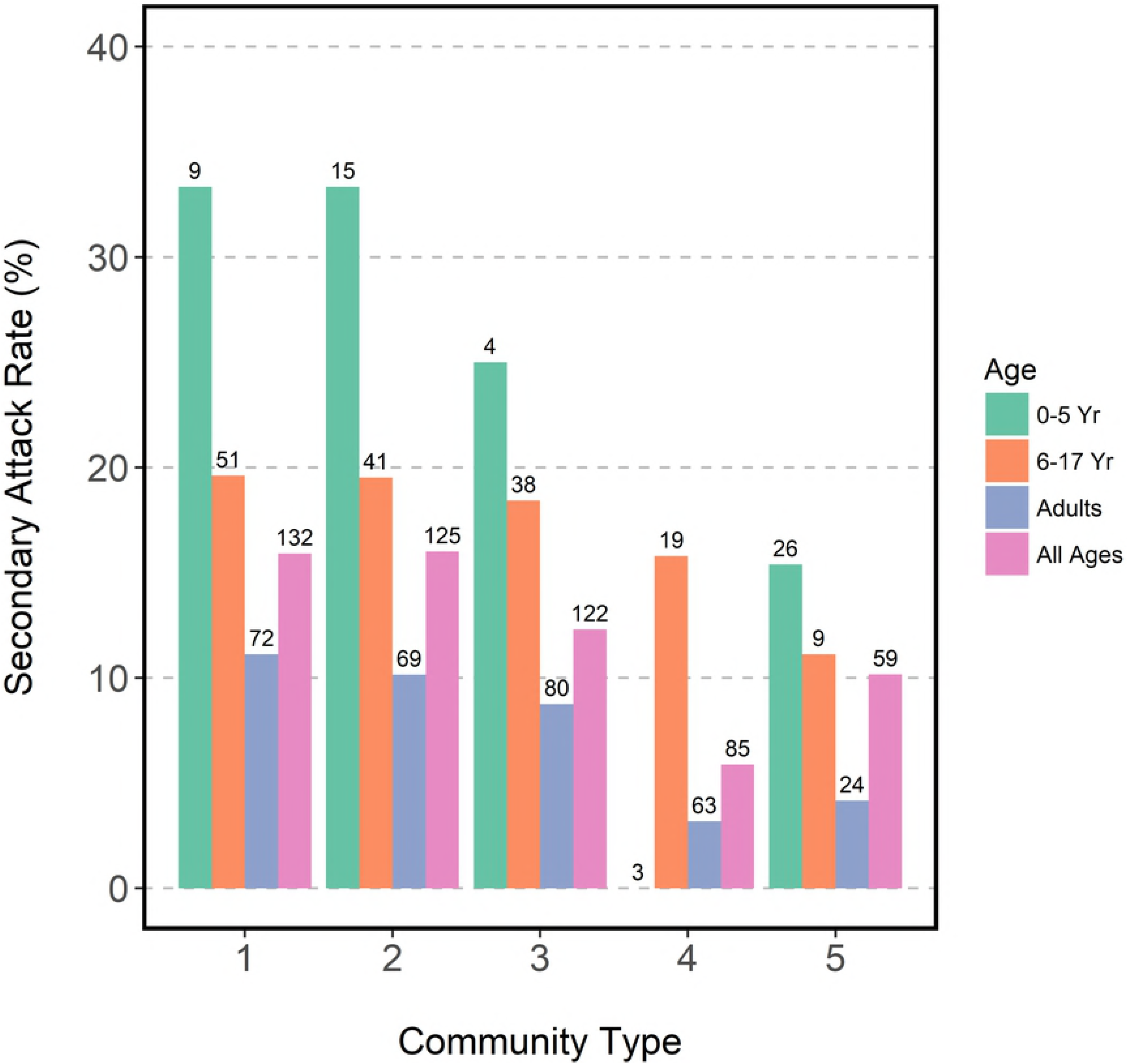
Secondary attack rates by community type at enrollment and age. 533 household contacts of influenza cases with defined community type at enrollment, residing in 144 households in Managua, Nicaragua, 2012-2014. Numbers represent sample size of each group.

**Fig 4.**
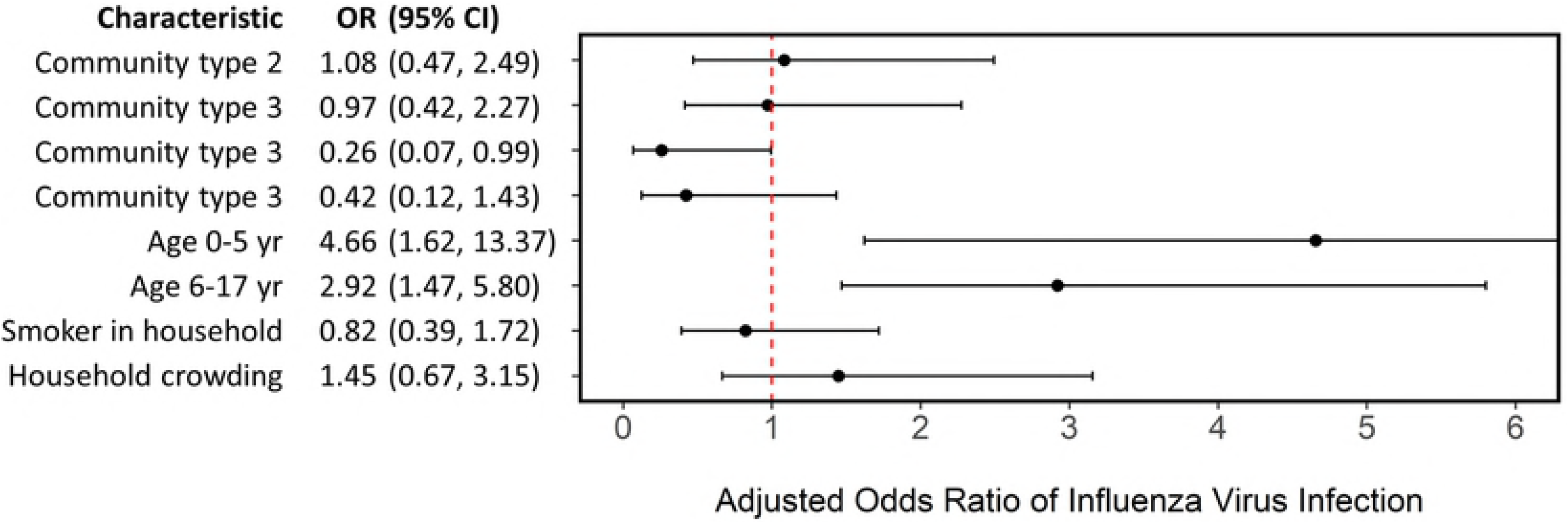
Generalized linear mixed effects model estimating odds of influenza virus infection. Model adjusting for community type (relative to community type 1), age (relative to adults), a smoker in the household, household crowding (average of >3 persons per bedroom), and clustering by household. 468 household contacts of influenza cases with complete data, residing in 132 households in Managua, Nicaragua, 2012-2014.

We were inadequately powered for influenza type/subtype-specific models; however, no household contacts with community type 4 at enrollment (n=85) were infected with H3N2, the most commonly identified influenza subtype in this population (52% of all secondary cases). This suggests associations between the microbiota and influenza susceptibility may vary by subtype but further work is needed to test this hypothesis.

### Oligotypes associated with susceptibility to influenza virus infection

In addition to analysis at the community type level, taxa-specific analysis was conducted using MaAsLin [27]. MaAsLin first uses boosting in a univariate pre-screen to identify taxa and metadata (features) that are potentially associated. Significantly associated features are then identified using linear mixed effects models. Models included in our analysis adjusted for age, a smoker in the household, household crowding, and clustering by household (S1 Appendix). Two oligotypes, *Alloprevotella sp.* and *Prevotella histicola / sp. / veroralis / fusca / scopos / melaninogenica,* were positively associated with influenza virus infection (S4 Table). One oligotype was negatively associated with influenza virus infection. Although unclassified in the HOMD database, a nblast search using the GenBank database (https://www.ncbi.nlm.nih.gov/genbank/) classified the oligotype as *Bacteroides vulgatus*.

The relative abundance of multiple oligotypes were strongly associated with age. Relative to adults, 119 oligotypes were differentially abundant among young children and 41 oligotypes were differentially abundant among older children. Lastly, four oligotypes were associated with household crowding and no oligotype was associated with exposure to a smoker in the household. All statistically significant associations are listed in S4 Table.

### Community diversity and influenza virus infection

We examined whether community diversity was associated with influenza susceptibility. Shannon diversity was significantly different between community type 4 and other community types (Wilcox rank-sum tests, all comparisons p<0.001) (S3 Fig). Community type 4 (median: 3.43) was less diverse than community type 1 (median: 3.58) and more diverse than community types 2, 3, and 5 (medians: 2.56-3.29).

To further explore whether community diversity influenced the relationship between community types and influenza susceptibility, we reran our generalized linear mixed effects model using Shannon diversity as our primary predictor. Alpha diversity was not significantly associated with influenza susceptibility (OR: 1.76; 95% CI: 0.83, 3.72) (S4 Fig).

### Stability of bacterial community structure during influenza virus infection

To examine the stability of the respiratory microbiota during influenza virus infection, we characterized changes in the bacterial community structure over a median of 9 days (IQR: 9, 10). We used Markov chain plots to represent short-term changes in the nose/throat microbiota among household contacts, by influenza case status and age (Fig 5 and S2 Appendix). Circles represent community types 1 through 5 and the size of the circles represent the prevalence of each community type at start of follow up. Arrows represent transitions between community types during follow up. The width and number assigned to each arrow represents the proportion of individuals who transitioned between community types. Community stability was estimated as the proportion of household contacts who showed no change in community type over follow up.

**Fig 5.**
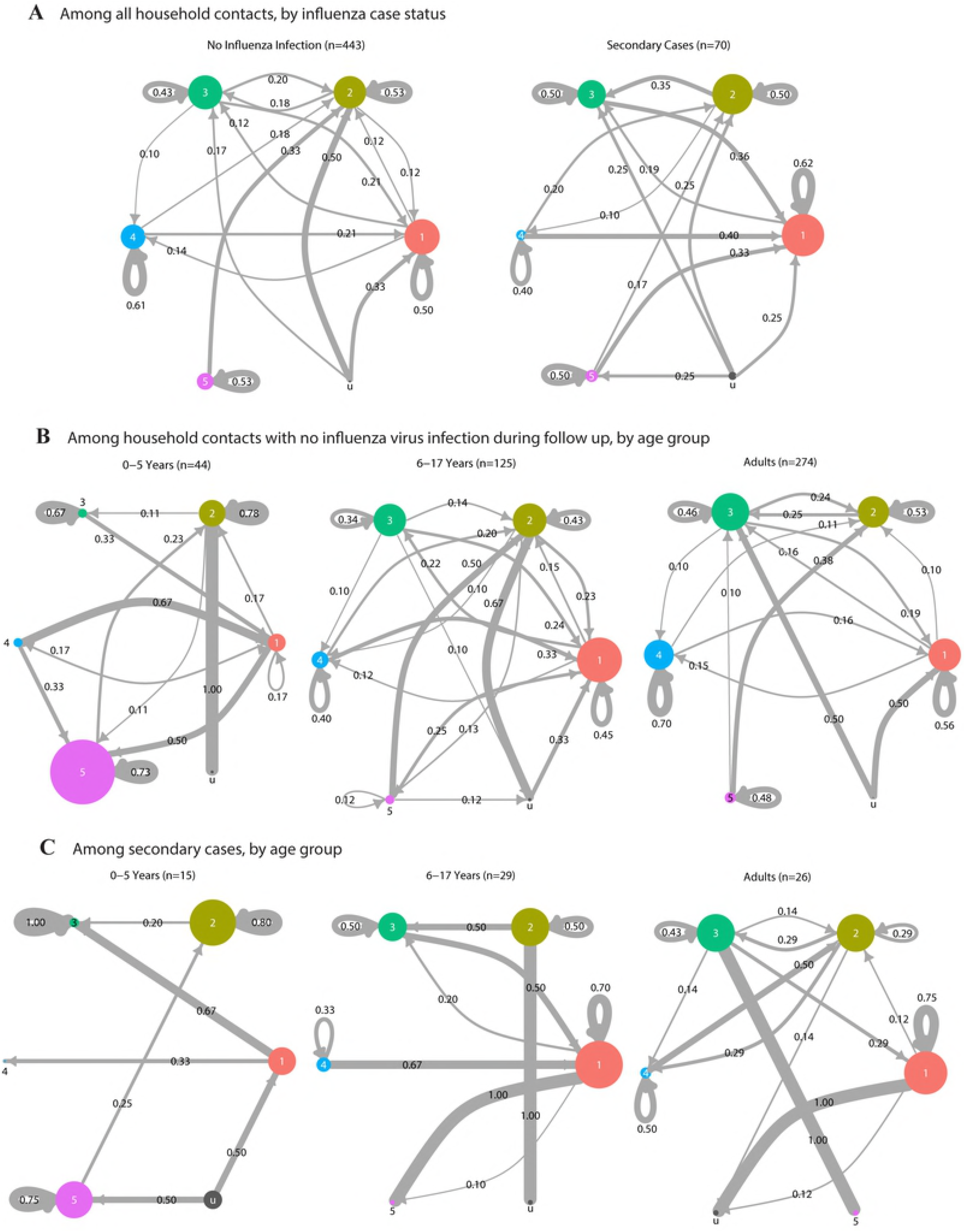
Stability of nose/throat bacterial community over follow up. 513 household contacts with microbiota data both at enrollment and follow up, residing in 144 households in Managua, Nicaragua, 2012-2014. **(A)** By influenza case status. **(B)** By age, among 443 household contacts who remained negative for influenza virus infection during follow up. **(C)** By age, among 70 secondary cases. Circles represent community types and circle size is proportional to prevalence of community types at enrollment. Community type u corresponds to samples with an undefined community type. Transition rates between community types were estimated as Markov chain probabilities and are shown numerically. Transitions rates <0.10 were removed for simplicity. Complete data are available in S2 Appendix.

Although the prevalence of community types appeared to remain similar between the two sampling points (S3 Table), we found transitions between community types were common with approximately half of all household contacts (45% among secondary cases, 55% among uninfected) changing to a different community type by the end of follow up (Fig 5 and S2 Appendix). Stability ranged from 40-62% for all community types in both secondary cases and uninfected household contacts. Although we were inadequately powered to test for statistical differences in specific type-to-type transitions, the overall contrast between the two groups suggest community dynamics may differ by influenza status and should be explored further.

We specifically focused on the stability of community type 4, which was associated with decreased influenza susceptibility. Stability among uninfected household contacts with community type 4 increased with age (0-5 years: 0%, 6-17 years: 40%, adults: 70%; Fisher exact test, p=0.016) (Fig 5B). We were inadequately powered for a similar analysis among secondary cases.

We used a generalized linear mixed effects model to examine whether community stability was associated with influenza virus infection, after adjusting for community type at enrollment, age, a smoker in the household, household crowding, and clustering by household (S1 Appendix). We did not find an association between community stability and influenza virus infection. However, we found stability was lowest among children 6-17 years old (OR: 1.67; 95% CI: 1.07, 2.60) (Fig 6).

**Fig 6.**
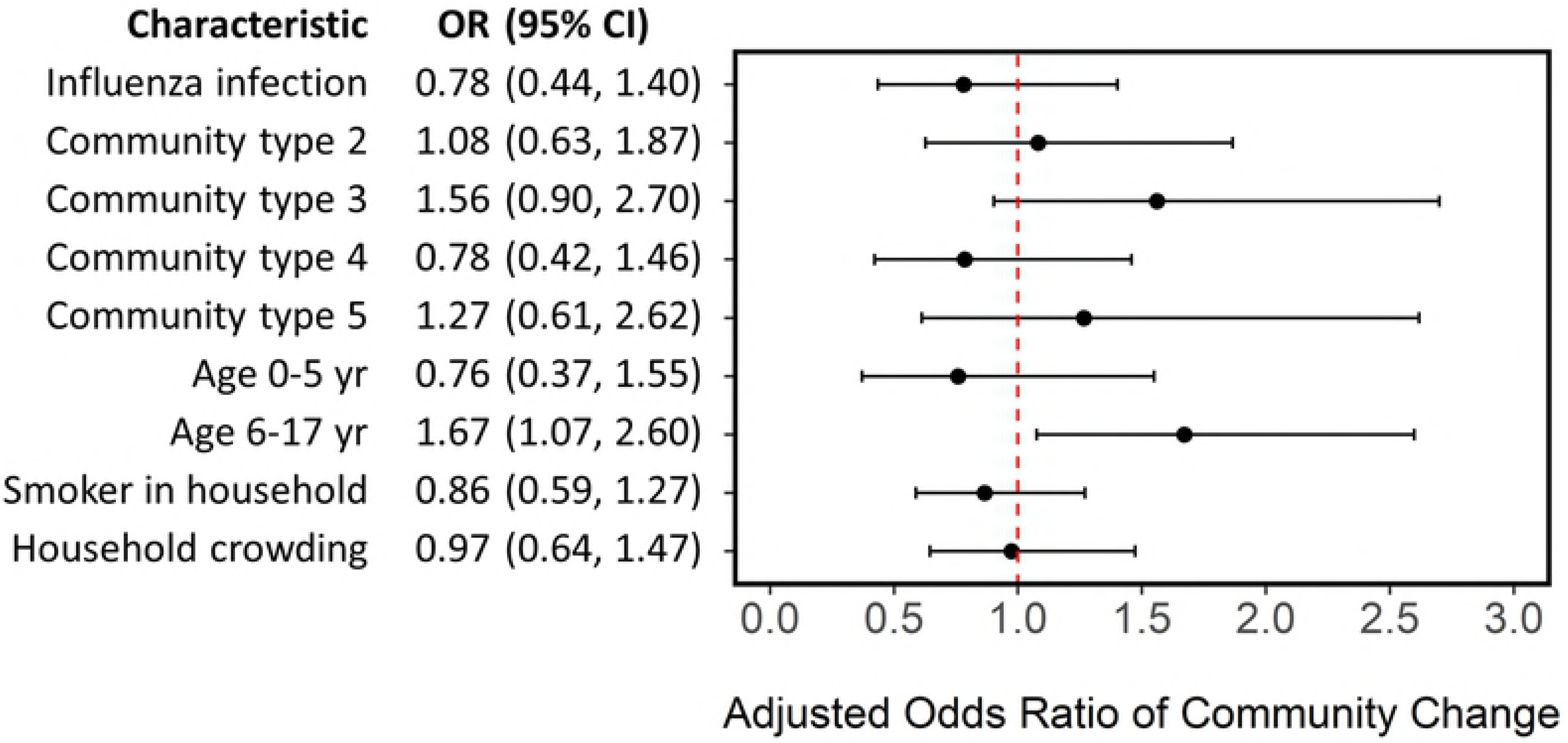
Generalized linear mixed effects model estimating odds of change in community type during follow up. Model adjusting for influenza virus infection, community type at enrollment (relative to community type 1), age (relative to adults), a smoker in the household, household crowding (average of >3 persons per bedroom), and clustering by household. 443 household contacts with defined community types at enrollment and follow up and complete data, residing in 130 households in Managua, Nicaragua, 2012-2014. Household contacts with an undefined community type were excluded from analysis.

### Sensitivity Analysis

Sensitivity analysis was conducted to investigate potential sources of bias (S3 Appendix). To assess whether sequencing depth could influence our results, we first examined whether sequencing depth differed by community type. We found no meaningful differences in sequencing depth by community type. In addition, we reran our influenza susceptibility model with sequencing depth as an additional predictor. We found only minor differences in model estimates, with a slightly enhanced effect of community type 4 (OR: 0.24; 95% CI: 0.06, 0.94).

To assess whether time between samples influenced community stability, we additionally controlled for time between samples in our community stability model. We found only minor differences in model estimates.

Lastly, we explored whether a more conservative criterion for community type assignments would influence our results. We reran our influenza susceptibility model after assigning samples with a maximum posterior probability ≤90% as missing. We found minor differences in our results. However, the association with community type 4 was no longer statistically significant (OR: 0.27; 95% CI: 0.07, 1.03).

## Discussion

To our knowledge, this is the first human population study to prospectively explore the relationship between the nose/throat microbiome and influenza virus infection. We demonstrate influenza susceptibility is associated with both differences in the overall bacterial community structure and in the relative abundance of specific taxa. Although the exact biological mechanisms remain unclear, the few murine studies that have examined this relationship suggest it is likely mediated by immunomodulation. In these studies, mice treated with antibiotics exhibited diminished innate and adaptive immune responses compared to placebo. Specifically, mice with disrupted microbiomes expressed impaired macrophage responses to type I and type II interferons and a lack of bacterial lipopolysaccharides that stimulate Toll-like receptors and other pattern recognition receptors [6,7]. Although these mechanisms suggest a causal relationship between the respiratory microbiome and influenza virus infection, additional work is needed to evaluate whether epidemiologic associations in human populations represent a true effect of the microbiome or merely reflect differences in host immunity. Further, future studies should examine whether the relationship differs by influenza type/subtype.

Most studies that have characterized the upper respiratory tract microbiota are limited to infants [22,28,29]. Here, we examine a unique population consisting of both children and adults. We demonstrate age is strongly associated with both the prevalence and stability of nose/throat bacterial communities. Most notably, we found the community type associated with decreased susceptibility to influenza was less prevalent and less stable among young children. If a causal relationship between the microbiome and influenza truly exists, our results would suggest the microbiome may contribute to the increased influenza risk observed among young children [30].

We found the microbiome structure changed frequently among both influenza cases and household contacts who remained uninfected during follow up (median: 9.0 days, IQR: 9.0-10.0). This was expected among influenza cases as prior studies have demonstrated increased colonization by opportunistic pathogens in the upper respiratory tract following respiratory virus infection [20,21]. However, the high degree of change among uninfected household contacts was surprising and may represent a response to influenza exposure in the household.

Preliminary findings from our Markov chain analysis suggest community dynamics may differ by influenza status. Characterizing multiple longitudinal samples per participant would lead to a better understand of influenza and its impact on the microbiome. In addition, future studies should explore whether changes in the bacterial community structure are directly due to influenza virus or indirect responses to changes in the virome or mycobiome.

A limitation in our study is the use of pooled nose and throat samples. Differential sampling by site can introduce bias if sampling is related to both the observed bacterial community structure and influenza susceptibility. Although this was minimized through consistent sampling techniques across all study participants, factors such as age could confound associations and may partially explain age-related differences in the bacterial community structure. We attempted to reduce potential bias by adjusting age and other potential confounders in our analysis. A second limitation is the use of RT-PCR for identifying influenza cases. Individuals can be infected with influenza virus (i.e. ≥4-fold increase in hemagglutinin inhibition antibody titer) and not shed virus [31]. We may have missed true index cases and secondary cases with low levels of virus. Although our results and conclusions are limited to secondary cases with viral shedding, RT-PCR allowed us to screen for secondary cases at 2-3 day intervals while limiting invasive procedures. Lastly, we did not consider pre-existing immunity from previous infections, which might potentially confound or modify associations.

While much work is needed to translate these results into potential clinical and public health applications, our findings contribute to a growing literature suggesting that it may be possible to manipulate the microbiome and decrease risk of disease [9,10]. Influenza virus is a major cause of severe illness and death each year [1,2]. However, vaccine effectiveness varies by year [4] and there still much debate on the use of antivirals for prophylaxis, especially for preventing asymptomatic infections and influenza transmission [32]. Our study suggests the microbiome should be further explored as a potential target in reducing influenza risk.

## Methods

### Study population and sample collection

The Nicaraguan Household Transmission Study of Influenza is an ongoing prospective case-ascertained study conducted among urban households in Managua, Nicaragua. Patients attending the Health Center Sócrates Flores Vivas were screened for study eligibility. Index cases of influenza were identified as patients with a positive QuickVue Influenza A+B rapid test, symptom onset of an acute respiratory infection within the past 48 hours, and living with at least one other household member. Symptoms of acute respiratory infection included fever or feverishness with cough, sore throat, or runny nose.

Index cases and household members (contacts) were invited to participate and clinical, sociodemographic, and household data were collected at time of enrollment. Participants were followed for up to 13 days through 5 home visits conducted at 2-3 day intervals. At each home visit, oropharyngeal and nasal swabs were collected, combined, and stored at 4°C in viral transport media. All samples were transported to the National Virology Laboratory at the Nicaraguan Ministry of Health within 48 hours of collection and stored at -80°C. A symptom diary was collected for all participants during follow up.

A total of 168 households were enrolled for follow up during 2012-2014. Households were excluded from analysis if a suspected index case was negative for influenza virus by real-time reverse-transcription polymerase chain reaction (RT-PCR) at time of enrollment. Two household contacts were excluded from analysis due to missing influenza virus infection status at time of enrollment. The remaining participants consisted of 144 index cases of influenza positive by RT-PCR, 537 household contacts influenza negative by RT-PCR at time of enrollment, and 36 household contacts who were RT-PCR positive for influenza virus on the first day of follow up.

### Ethics statement

Written informed consent was obtained from adult participants and from parents or legal guardians of participants under 18 years of age. In addition, verbal assent was obtained from children over 5 years of age. The study was approved by Institutional Review Boards at the University of Michigan, the Nicaraguan Ministry of Health, and the University of California at Berkeley.

### RNA extraction and RT-PCR

Total RNA was extracted from all available nasal/oropharyngeal samples using the QIAmp Viral Mini Kit (QIAGEN, Hilden, Germany) per manufacturer’s instructions at the National Virology Laboratory in Nicaragua. Samples were tested for influenza virus by RT-PCR using standard protocols validated by the Centers for Disease Control and Prevention [33].

### DNA extraction and 16S rRNA sequencing

Total DNA was extracted from a pair of samples from each study participant: the first sample collected at time of enrollment and the second sample collected at the last day of follow up (median days between samples: 9.0 days, IQR: 9.0-10.0). Among the 717 total study participants, five first samples and 19 second samples were not available for DNA extraction. DNA was extracted using the QIAmp DNA Mini Kit and an enzyme cocktail composed of cell lysis solution (Promega, Madison, USA), lysozyme, mutanolysin, RNase A, and lysostaphin (Sigma-Aldrich, St. Lious, USA) in 22.5:4.5:1.125:1.125:1 parts, respectively. 100 μL of sample was incubated at 37°C for 30 minutes with 80 μL of the enzyme cocktail. After adding 25 μL proteinase K and 200 μL of Buffer AL, samples were vortexed and incubated at 56°C for 30 minutes. Samples were washed with 200 μL of 100% ethanol, 500 uL of Buffer AW1, and then 500 uL of Buffer AW2. To maximize DNA yield, DNA was eluted twice with 100 uL of Buffer AE and stored at - 80°C.

The V4 hypervariable region of the 16S rRNA gene was amplified and sequenced at the University of Michigan Microbial Systems Laboratories using Illumina MiSeq V2 chemistry 2×250 (Illumina, San Diego, CA) and a validated dual-indexing method [34]. Briefly, primers consisted of an Illumina adapter, an 8-nt index sequence, a 10-nt pad sequence, a 2-nt linker, and the V4-specific F515/R806 primer [35]. Amplicons were purified and pooled in equimolar concentrations. A mock community of 21 species (Catalog No. HM-782D, BEI Resources, Manassas, VA) or a mock community of 10 species (Catalog No. D6300, Zymo Research, Irvine, CA) was included by the Microbial Systems Laboratories to assess sequencing error rates. For every 96-well plate submitted for amplification and sequencing (90 study samples), we included two aliquots of an in-house mock community consisting of *Streptococcus pneumoniae, Streptococcus pyogenes*, *Staphylococcus aureus, Haemophilus influenzae,* and *Moraxella catarrhalis* and two aliquots of an oropharyngeal control sample. These internal controls were randomly assigned to plate wells and used to assess systematic variation in sequencing. All samples were sequenced in duplicate, demultiplexed, and quality filtered.

### Oligotyping and community typing

We used mothur v1.38.1 [36] to align and perform quality filtering on raw sequences using the mothur standard operating procedures (https://www.mothur.org/wiki/MiSeq_SOP, accessed November 18, 2016). Sequences were converted to the appropriate oligotyping format as previously described [37]. We used the Minimum Entropy Decomposition (MED) algorithm [38] with default parameters (-M: 13779.0, -V: 3 nt) to cluster sequences into oligotypes. Briefly, the algorithm identifies variable nucleotide positions and uses Shannon entropy to partition sequences into nodes. The process is iterative and continues to decompose parent nodes into child nodes until there are no discernable entropy peaks. Oligotyping has previously been used to examine within-genus variations in the microbiota [37, 39–41] and provides increased resolution relative to conventional distance-based clustering methods.

After excluding five samples with less than 1,000 reads, our dataset consisted of 1,405 samples with a total of 61,784,957 sequences decomposed into 230 oligotypes. To assign taxonomy, we searched representative sequences of each oligotype against the Human Oral Microbiome Database (HOMD) v14.51 [42] using blastn v2.2.23 [43]. Classifications with ≥98% identity were kept.

We used Dirichlet multinomial mixture models [26] in R v3.4.4 [44] and the DirichletMultinomial v1.16.0 package [45] to assign all samples to 5 community types. We determined the number of community types by comparing the Laplace approximation of the negative log models and identifying the point at which an increase in Dirichlet components resulted in minor reductions in model fit (S1 Fig). This approach allowed us to consider both model fit of the negative log models and statistical power in downstream analysis. The goal was not to identify the “true communities”, as community types are representations of data. All formal statistical inferences are based on the models relating community types to our outcomes of interest, with any findings being statistically supported by the data.

Samples were assigned to community types with the greatest posterior probability. 98.2% of all samples had a posterior probability of 80% or higher. To minimize misclassification, samples were assigned as having an undefined community type if the posterior probability was less than 80%. Each community type contained between 13.0-24.8% of all samples (n=182-348) and 1.8% of all samples (n=25) were undefined.

### Statistical Models

Detail to statistical models used in this study are described in S1 Appendix. To examine the association between community types at enrollment and susceptibility of influenza virus infection, we used a generalized linear mixed effects model estimating the odds of infection after adjusting for community type (relative to community type 1), age (relative to adults), a smoker in the household, household crowding, and clustering by household. Household crowding was defined as having, on average, more than three household members per bedroom. The model was adapted to examine the effects of alpha diversity.

Associations between individual oligotypes and participant data were determined using MaAsLin [27]. Briefly, MaAsLin is a sparse multivariate approach used to identify associations between individual taxa and participant data. Relative abundance values were arcsine square-root transformed to stabilize variance. Potentially associated features (i.e. oligotypes) were selected using boosting in a univariate prescreen. Linear mixed effects models are then used to find associations between the selected features and metadata. The Benjamin-Hochberg method was used to correct for multiple testing. Associations with a q-value <0.05 were considered statistically significant.

To examine the effect of influenza virus infection on community stability, we used a generalized linear mixed effects model estimating the odds of any change in community type over follow up, after adjusting for community type at enrollment (relative to community type 1), age (relative to adults), a smoker in the household, household crowding (average of >3 persons per bedroom), and clustering by household. All statistical analysis was conducted using R v3.4.4 [44] and the lme4, vegan, and maaslin packages [27,46,47].

### Markov chain analysis

We estimated community transition rates over time using methods described previously [48]. Briefly, we restricted our dataset to household contacts with complete nose/throat sample pairs (i.e. microbiota data at enrollment and at follow up). Community transition rates were calculated as Markov chain probabilities. Analysis was repeated after stratifying by influenza status and age.

## Acknowledgments

We are grateful to the participants and our collaborators at the Nicaraguan Ministry of Health and Sustainable Sciences Institute for data collection, data management, and laboratory work. We thank Roger Lopez for conducting RT-PCR. We also thank the University of Michigan Microbial Systems Laboratories for 16S sequencing and Nielson Baxter, Michelle Berry, and Ruben Props for technical assistance in mothur and oligotyping. We thank Anna Gencay and Sarah McColm for laboratory assistance. We appreciate feedback we received throughout the course of the study from Carl Marrs, Alex Rickard, Rachel Gicquelais, Kirtana Ramadugu, Freida Bolstein, Emily Menard, and our MAC-EPID colleagues. We thank Marie Griffin and Jae Shin for reviewing an earlier version of this manuscript. We are entirely grateful for all the study participants in the Nicaraguan Transmission Study of Influenza.

---

**S1 Fig. Model fit of negative log models by number of Dirichlet components.**

We determined the number of community types by estimating the Laplace approximation of the negative log models and identifying the point at which an increase in Dirichlet components resulted in minor reductions in model fit. This approach allowed us to consider both model fit of the negative log models and statistical power in downstream analysis. The goal was not to identify the “true communities”, as community types are representations of data. All formal statistical inferences are based on the models relating community types to our outcomes of interest, with any findings being statistically supported by the data.

**S2 Fig. Principal coordinates analysis of nose/throat samples assigned to community types.**

1,405 nose/throat samples from 717 study participants residing in 144 households in Managua, Nicaragua, 2012-2014. Based on Bray-Curtis dissimilarity.

**S3 Fig. Shannon diversity of nose/throat samples, by bacterial community type.**

1,380 samples with defined community type, 717 study participants residing in 144 households in Managua, Nicaragua, 2012-2014.

**S4 Fig. Generalized linear mixed effects model estimating odds of influenza virus infection using Shannon diversity**.

Model adjusts for Shannon diversity, age (relative to adults), a smoker in the household, household crowding (average of >3 persons per bedroom), and clustering by household. 477 household contacts of influenza cases with complete data, residing in 132 households in Managua, Nicaragua, 2012-2014.

**S1 Table. Characteristics of 71 secondary cases from 48 households, Managua, Nicaragua, 2012-2014, by community type at enrollment.**

**S2 Table. Relative abundance of all 230 oligotypes, by community type.**

A total of 1,405 samples from 717 study participants residing in 144 households in Managua, Nicaragua, 2012-2014.

**S3 Table. Distribution of community types by age, time, and whether acquired influenza by end of follow up.**

537 household contacts with any microbiota data, residing in 144 households in Managua, Nicaragua, 2012-2014.

**S4 Table. MaAsLin results.**

**S1 Appendix. Description of statistical models.**

**S2 Appendix. Stability of nose/throat bacterial community over follow up, including all transitions.**

**S3 Appendix. Sensitivity analysis.**

